# Neural networks and extreme gradient boosting predict multiple thresholds and trajectories of microbial biodiversity responses due to browning

**DOI:** 10.1101/2021.03.22.435765

**Authors:** Laurent Fontaine, Maryia Khomich, Tom Andersen, Dag O. Hessen, Serena Rasconi, Marie L. Davey, Alexander Eiler

## Abstract

Ecological association studies often assume monotonicity such as between biodiversity and environmental properties although there is growing evidence that non-monotonic relations dominate in nature. Here we apply machine learning algorithms to reveal the non-monotonic association between microbial diversity and an anthropogenic induced large scale change, the browning of freshwaters, along a longitudinal gradient covering 70 boreal lakes in Scandinavia. Measures of bacterial richness and evenness (alpha diversity) showed non-monotonic trends in relation to environmental gradients, peaking at intermediate levels of browning. Depending on the statistical methods, variables indicative for browning could explain 5% of the variance in bacterial community composition (beta diversity) when applying standard methods assuming monotonic relations and up to 45 % with machine learning methods (i.e. extreme gradient boosting and feed-forward neural networks) taking non-monotonicity into account. This non-monotonicity observed at the community level was explained by the complex interchangeable nature of individual taxa responses as shown by a high degree of non-monotonic responses of individual bacterial sequence variants to browning. Furthermore, the non-monotonic models provide the position of thresholds and predict alternative bacterial diversity trajectories in boreal freshwater as a result of ongoing climate and land use changes, which in turn will affect entire ecosystem metabolism and likely greenhouse gas production.

## Introduction

For simplification, ecological associations such as between biodiversity and environmental properties are often assumed to be monotonic, i.e. either positive, negative or neutral. But in nature, non-monotonic interactions are commonly seen at the individual, population, community and ecosystem levels. Most non-monotonic relations reported in the ecological literature are periodic cycles in time (i.e. prey and predator relationship, ref. [1]) or humped-shaped curves when inferring for example relationships between productivity and biodiversity [2–4]. Non-monotonicity has been suggested to represent an important driving force in ecological systems because environmental factors are highly variable in both space and time, and organisms do not interact with abiotic and biotic factors in a fixed way [5]. A common feature of non-monotonic functions is that they define relationships with both increasing and decreasing sectors as well as different stable states where the nature of the response can change dramatically when an environmental factor (i.e. temperature) reaches a threshold (or ridge). Such thresholds are missed by monotonic (linear) models commonly used in ecological data interpretation and modeling. The assumption of monotonicity and resulting over-simplification of biological complexity has been criticized by many ecologists [6, 7].

An intensively studied relationship in microbial ecology is the link between microbial diversity and natural organic matter (NOM) which represents a major energy source for heterotrophic bacteria [8]. By far the largest NOM pool in aquatic environments is dissolved organic matter (DOM) which is of a complex and heterogenous nature [9]. Subsets of the diverse DOM pool can have a strong influence on light attenuation, metal speciation, and bioavailability, while also acting as a pH buffer [10]. In recent decades an increase of DOM loadings to boreal surface waters has been observed [11, 12]. This increase has been linked to a 30 % increase of precipitation due to climate change and a projected 15-20% increase in runoff [13]. Exacerbated by land use change, the increased supply of DOM to lakes and rivers [14] has direct and indirect effects on the microbial loop with implications for phenological events such as timing of the spring phytoplankton bloom [15] and fish spawning time. Also, increased levels of chromophoric DOM will suppress primary production due to light limitation [16], while providing substratum for heterotrophic bacteria [8, 17], thereby promoting reduced production to respiration ratios. Thus, overall changes to carbon processing by heterotrophic bacterial communities can affect emissions of CO2 and CH4 from the boreal landscape and local water quality [18–20].

Complex interactions between heterotrophic bacteria and DOM have been suggested to shape the apparent composition of both of these key ecosystem components [21–25]. This coupling is corroborated by incubation experiments under controlled laboratory conditions where it has been shown that the availability and composition of organic substrates favor specific bacterial groups, and in this way shape bacterial community composition (BCC) and community metabolism [26–30]. Moreover, bacteria do not only consume and degrade DOM, but also produce and release an array of autochthonous organic compounds during cell growth, division, and death [31], thereby influencing the availability, composition, and biogeochemical cycling of C in the biosphere [32, 33]. While community adaptation (i.e. composition shifts) has been found to precede bacterial degradation of specific carbon substrates [34], the contribution of bacterial community shifts and key bacterial players to the production and degradation of DOM is unclear [5]. As a result of these multiple levels of interactions and feedbacks, relationships between DOM and bacterial diversity are expected to be non-monotonic.

Our study is based on samples from 70 large and relatively deep boreal lakes along a 750 km longitudinal gradient across southern Scandinavia. The Scandinavian diversity gradient is complex and not fully resolved as it coincides both with the main post-glacial dispersal routes for freshwater biota, as well as with major changes in soil depth, altitude and landscape productivity [35]. Previous molecular [35, 36] and non-molecular [37] studies have described the diversity and community composition of pelagic protists, aquatic fungi, zooplankton and fish along a longitudinal gradient in these lakes. Generally, there is a strong decline in diversity across functional and taxonomic groups from east to west. The survey covers a wide longitudinal range and broad gradients in DOM quality and quantity as well as nutrient status of the systems allowing us to parse out the spatial vs. local environmental effects on bacterial biodiversity.

Here we aim to capture non-monotonic features by using modern statistical tools such as generalized-additive-models, maximal-information-based-nonparametric-exploration (MINE), marginal-(maximum)-likelihood-model-fitting, eXtreme-Gradient-Boosting (XGBoost) and feed-forward-neural-networks (FFNN). We tested the hypothesis that threshold responses and alternative trajectories exist in biodiversity responses due to browning in freshwater lakes. Taking into account co-varying factors such as nutrient status and other environmental abiotic gradients, we trained XGBoost and FFNNs to predict the interactions between DOM and bacterial community composition in the studied systems so as to identify thresholds in community composition along the studied DOM gradient. Ultimately, we intend to interpolate our findings in the light of ongoing environmental change.

## Materials and methods

### Site description and sampling

Lakes were selected from the ‘Rebecca’ [38] and ‘Nordic lake survey 1995’ [39] data sets on Norwegian and Swedish lakes to create a subset fulfilling the following criteria: longitude 5 – 18 °E, latitude 58 – 62 °N, altitude < 600 m, surface area > 1 km^2^, total phosphorus (TP) < 30 μg L^-1^, total organic carbon (TOC) < 30 mg L^-1^ and pH > 5. Acidic, eutrophic and highly dystrophic lakes were omitted. The final subset represents similarly sized boreal lakes within a narrow latitudinal and altitudinal range, with the best possible coverage and a tentative orthogonality with respect to gradients of TP, TOC and longitudinal position. In particular, longitude reflects the regional diversity gradient described in ref [40], while TP and TOC represent two major and contradictory effects on aquatic productivity [16]. Water temperature, pH and conductivity were measured *in situ*, and samples for nutrient analysis were collected as described in ref. [16]. There is a strong relationship between snap-shot temperature measured with the CTD and climatic average mean July air temperature, suggesting that the longitudinal temperature gradient is not confounded by the sampling scheme starting the survey in the west and moving eastward across the gradient. At each site, a water sample was collected from the lake epilimnion (0-5 m) in the central part of each lake during daytime using an integrating water sampler (Hydro-BIOS, Germany). For DNA extraction, up to 100 mL of water was pre-screened *in situ* on 100 µm mesh to remove large non-microbial cells and then filtered through 0.2 μm pore size polycarbonate filters (25 mm diameter; Poretics, Spectrum Chemical Corp., NJ, USA) taken in 3 replicates. The filters were frozen in liquid nitrogen *in situ* and subsequently stored at -20°C in cryovials until DNA extraction. The detailed sampling strategy and analytical methods have been previously described [16, 35, 36].

### Carbon characterization

TOC was measured by infrared CO2 detection after catalytic high temperature combustion (using either a Shimadzu TOC-VWP analyzer or Phoenix 8000 TOC-TC analyzer). Particulate organic carbon (POC) was measured on an elemental analyzer (Flash EA 1112 NC, Thermo Fisher Scientific, Waltham, Massachusetts, USA) through rapid combustion of a pre-combusted GF/C filter with particulates in pure oxygen, where carbon was detected as CO2 by gas-chromatography. DOC was calculated as the difference between TOC and POC. Carbon quality was assessed via absorbance spectra. After lake water had been filtered through a Acrodisc 0.2 µm polyethersulfone membrane syringe filter (Pall Life Sciences), the optical density of the filtrate (OD_CDOM_(λ)) was measured in a 50-mm glass cuvette from 400 to 750 nm in steps of 1 nm. Absorption coefficient spectra of chromophoric DOM (αCDOM(λ); m− 1) were calculated according to ref. [41].

The absorbance measured at 400 nm (a_CDOM_(400)) was used as a proxy for aromaticity of chromophoric DOM (CDOM) after dividing by TOC concentrations. Iron can bind to humic substances and form complexes that may increase absorbance [42]. To account for this, a correction factor developed for a_CDOM_(400) using concentrations of dissolved iron Fe^3+^ was applied. Non-algal particulate matter (NAP) was assessed by the optical density (ODNAP(λ)), as described in ref. [16]. Absorption coefficients (m− 1) of total particulate matter (α_p_(λ)), and NAP (αNAP(λ)), were calculated according to ref. [41]. We used the algorithm of Bricaud and Stramski [43] to estimate the path-length amplification factor (β). Finally, we calculated the absorption coefficient spectra of phytoplankton pigments (α_ph_(λ); m− 1) as the difference between the total particulate and the NAP absorption coefficient spectra.

### DNA extraction, amplification and Illumina HiSeq sequencing of the V4 SSU

Total DNA was extracted from the filters using the PowerSoil DNA isolation Kit (MoBio Laboratories Inc., Carlsbad CA, USA) according to the manufacturer’s instructions and quantified using Qubit 2.0 Fluorometer (Invitrogen). The extracted DNA was sent to GATC Biotech (Konstanz, Germany) for amplification and HTS amplicon sequencing (INVIEW Microbiome Profiling 2.0 package). A set of universal primers was used to amplify the hypervariable regions V3-V5 (∼569 bp) of the 16S rRNA gene. Amplicon sequencing was done on an Illumina HiSeq Rapid Run instrument using a paired-end 300bp sequence run. The raw reads with corresponding mapping files were deposited in SRA under accession number PRJNA637765.

### Bioinformatics

Raw sequence data was processed with CUTADAPT [44] to remove primers and then analyzed using DADA2 [45]. Forward and reverse reads were trimmed at 200bp and 160bp, respectively. Reads were denoised using the DADA2 machine-learning algorithm. Since trimming resulted in no overlap of the read pairs, forward and reverse reads were concatenated. Quality filtering removed any paired reads with missing primers or ambiguous base pairs as well as a phred score below 20 somewhere in the paired reads. Taxonomic annotation was performed against the SILVA 132 database [46] using the Naive Bayesian classifier [47].

### Statistics

All downstream statistical analyses were performed in R version 3.6.0 [48] using vegan [49], PHYLOSEQ [50] and MASS [51] for multivariate and species richness analyses unless otherwise noted. Missing values in the metadata were approximated using multiple imputation with Fully Conditional Specification (FCS) implemented by the MICE algorithm as described in ref. [52].

CDOM variables used in this study included absorption coefficients at 400nm (a_CDOM_(400)) and absorption spectral data between 400-750 nm. The entire absorption spectral data was scaled and a principal component analysis (PCA) was performed resulting in a PCA model with principal component 1 (PC1) explaining over 88% of the variance. As such PC1 scores can be used as an index to characterize the CDOM variability among the samples. Partial least square modelling was performed with packages mdatools (function *randtest*) and plsdepot (functions *plsreg1* and *plsreg2* with cross-validation) using the first six principal components of the PCA from absorption data (Y variables), and scaled environmental data (X variables).

The two technical replicates were excluded from further downstream analyses as within replicate sequence variants were significantly more similar than between sample comparisons (data not shown). To calculate diversity measures, the sequence variant table was rarefied to a common sampling depth of 392 082 reads/sample, based on the sample with the least number of reads. Species accumulation curves (SAC; calculated using the analytical version of the *specaccum* function) were applied to assess sampling effort in each lake. Rarefaction curves were constructed for each lake using the *rarecurve* function in vegan. Alpha diversity indices (observed richness, Shannon and Simpson diversity) were calculated for each lake using the function *diversity* (R package vegan). Associations between alpha diversity indices and DOM descriptors were explored with generalized additive models (GAMs) using R package mgcv.

Non-metric multidimensional scaling (NMDS) ordinations [53] from multiple starting points (*metaMDS* function in vegan, *try* = 1000) were used to describe patterns in bacterial community composition (based on Hellinger-transformation and Bray-Curtis distance measure) between lakes. Permutation-based significance tests (*n* = 999) with the *envfit* function were used to fit spatial and environmental gradient variables to the NMDS ordination. The local environment was defined by the concentrations of total, particulate and dissolved CNP and other parameters (see Supplementary Table S1 for complete list of variables), while the spatial factors were represented by longitude, latitude and altitude. In addition, a redundancy analysis (RDA) was performed on bacterial community composition (Hellinger-transformed) using scaled environmental data.

To determine the relative role of DOM descriptors (a_CDOM_(400), PC1-CDOM and TOC), local (all other environmental variables) and regional (spatial factors) predictors on the distribution of bacterial communities along the biodiversity gradient, variance partitioning analysis was used. Variance partitioning by RDA (function *varpart* in vegan) [49] on Hellinger transformed, normalized abundance data was used to estimate the variance fractions of bacterial community composition that could be explained independently by the local environment divided into DOM and other parameters, spatial gradients (latitude and longitude), or shared between them. Marginal (Maximum) Likelihood model fitting was used to fit a smooth response surface of TOC and a_CDOM_(400) values over the limits of the biplot of the bacterial community composition using the *ordisurf* function.

Machine learning algorithms were used to identify beta diversity patterns along the CDOM and TOC concentration gradients. Regression was performed with a_CDOM_(400) and TOC values as inputs and BCC Bray-Curtis distances as outputs using scikit-learn’s implementation of XGBoost and random forest regressor (of the PyPi packages) as well as TensorFlow for the FFNNs using backpropagation [54]. In short, data was split in training and test sets comprising 80 and 20 % of observations, respectively. For the FFNN, the weights of hidden layers were initialized using Xavieŕs initialization [55], with ReLu activation and mean-squared error being used as a cost function. For visualization of the models, the original meshgrid of a_CDOM_(400) and TOC values spanning the minimum-maximum range of said gradient with a step size equal to the smallest pairwise a_CDOM_(400) and TOC differences. XGBoost stands for Extreme Gradient Boosting and represents a specific implementation of the Gradient Boosting method and uses more accurate approximations to find the best (decision-)tree model. In prediction problems involving unstructured data (images, text, etc.) neural networks tend to outperform all other algorithms or frameworks. However, when it comes to small-to-medium structured/tabular data as in our case, decision tree based algorithms are considered to be better suited. XGBoost is exceptionally successful, particularly with structured data since it computes second-order gradients, i.e. second partial derivatives of the loss function (similar to Newton’s method), which provides more information about the direction of gradients and how to identify minimae in the loss function. XGBoost uses the 2nd order derivative as an approximation and advanced regularization, which improves model generalization. Accuracy of the methods were compared using mean squared error (MSE) while the variance of raw data explained by the model was computed with R^2^.

In addition we performed ordinary least squares (OLS) regression by singular value decomposition (SVD) using polynomials of a_CDOM_(400) and TOC values as inputs. An appropriate polynomial degree was chosen in light of the bias-variance trade-off, where error was minimal while bias and variance curves intersected.

In the maximal information-based nonparametric exploration (MINE, ref. [56]) analysis run with default settings, relationships with p values of <0.05 were recorded with a false discovery rate, as determined by Hochberg, of <0.05 (q-values). The chosen p-value set the maximal information coefficient (MIC) cutoff to 0.3. The MIC is a statistical measure, similar to R^2^ in general linear models, describing the goodness of fit between two variables [56]. Various statistics can be used to characterize the relationships identified by MIC including measures of monotonicity, non-linearity, closeness to being a function, and complexity of relationships. The Maximum Asymmetry Score (MAS) measures deviation from monotonicity. We plotted the variability in MIC and MAS between amplicon sequence variants (ASVs) and a_CDOM_(400) or TOC with q-values < 0.05 and linked them with the sign of the correlation coefficient (Spearman R).

## Results

### The lake gradient through DOM quantity and quality

The sampled lakes spanned from the Norwegian Coast of the North Sea to the Swedish East Coast of the Baltic Sea and represent summer conditions as samples were taken from 20 July to 16 August 2011 (Figure 1A). Besides varying in latitude (58 – 62 °N) and temperature (9.9 – 21.4 °C), lakes varied in nutrient content with TOC in the lakes ranging from 0.3 to 12.9 mg l^-1^ (median 6.5 mg l^-1^), TP from 0.5 to 27.5 µg l^-1^ (median 4.55 µg l^-1^), total organic nitrogen (TON) from 87 to 1526 µg l^-1^ (median 298 µg l^-1^), and chlorophyll a from 0.77 to 29.5 (median 2.7). Lake size varied from 0.1 to 14 km^2^ with a median of 0.3 km^2^ (Table S1; ref. [16]).

**Figure 1.**
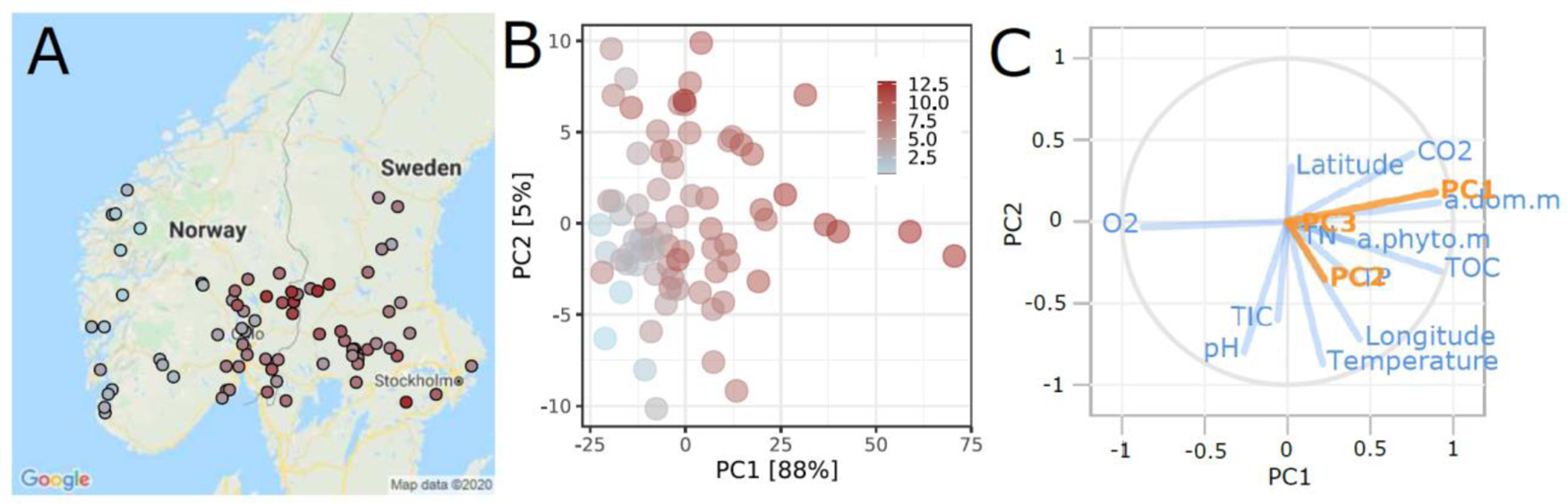
Map of sampling locations **(A)** with total organic carbon concentrations (mg L^-1^) in the lake system indicated by point colour. Principal component analysis (PCA) **(B)** for the quality of dissolved organic matter (DOM) as assessed by fluorescence spectroscopy. Partial least square (PLS) loading plot **(C)** revealing the covariation of the first three principal components for the quality of dissolved organic matter (CDOM) which were taken from analysis in panel B (Y variables in orange), and geographical, physical and chemical lake characteristics as predictors (X variables in blue). The comparison of observed and model predictions of CDOM are shown in supplementary figure S2 corroborating the high predictive power of the PLS model (R^2^ = 0.815 and Q^2^ = 0.775) when using environmental properties.

The PCA revealed substantial differences in DOM quality along the sampled lakes as assessed by absorption spectra (Figure 1B). Firstly, the relative positioning of the sample scores was mainly a function of PC1 which explained 88% of the variability. This component was a function of TOC concentration and a_CDOM_(400) as revealed by Spearman rank correlation analyses (R = 0.75; p < 0.0001 and R = 0.8; p < 0.0001, respectively). Other significantly correlated (p < 0.0001) environmental variables with CDOM (PC1) and a_CDOM_(400) included gas concentrations (O2, CO2, CH4), chlorophyll-a, total (TP) and particulate nutrient concentrations (PON, POC and POP) (see also Supplementary Figure S1).

Furthermore, partial least squares (PLS) was applied to predict CDOM (PC1-3) variability, (Y response variables) from lake water chemistry and climate variables (X predictor variables). Variables (X or Y) situated close together on the PLS plot such as CO2, TOC and a_CDOM_(400) can be interpreted as positively correlated with CDOM PC1, while variables opposite to CDOM PC1 as negatively correlated, such as O2 (Figure 1C). Next, we performed a PLS using only CDOM PC1 (Supplementary Figure S2A) with internal cross-validation to test the repeatability of the analysis by removing a random subset of data (1/7th of the samples) to be used as the response dataset, while parallel models were run on the reduced calibration dataset. A comparison of predicted values from the calibration and response datasets allowed computation of the predictive residual sum of squares, expressed as a Q^2^Y. Overall, PLS model performance was good with a cumulative goodness of fit (R^2^Y, explained variation) of 0.815, and the cumulative goodness of prediction (or Q^2^Y, predicted variation) of 0.775 for the PLS model with two components. This was also corroborated by comparing original and predicted values (Supplementary Figure S2B). As such the PLS model corroborates the results of the multiple correlation analysis (Supplementary Figure S1) with CO2, TOC and O2 concentrations representing environmental properties highly related to CDOM characteristics expressed by CDOM PC1 and a_CDOM_(400).

### Overall bacterial community features and diversity

A total of 15120 unique sequence variants (including 864 archaeal and eukaryotic reads) were recovered from 25 574 631 high-quality reads across 72 lakes. After removing non-bacterial reads, an average of 764 (range = 354–1454, SD = 208) ASVs were detected per sample and the mean number of reads per lake was 352 572 (range = 234 930–502 704, SD = 57 780). A total of 174 ASVs were detected in more than 50 lakes with the mean number of total reads per lake ranging from 219 to 5754 reads and representing some of the most abundant sequence variants in our dataset. Rarefaction curves of ASV richness (Supplementary Figure S3A) for each lake indicated that the total bacterial diversity was almost entirely recovered in all samples, since the rarefaction curves approached asymptote and sampling saturation. Still, region-wide species accumulation curves based on the progressive or random addition of samples showed that the gamma diversity in the studied area has not been fully recovered (Supplementary Figure S3B).

Various diversity indexes were highly correlated in the present dataset. For example, bacterial diversities calculated using inverse Simpson, Shannon, Fisher and ACE (abundance-based coverage estimators) diversity were highly correlated: R > 0.46, p < 0.0005. For example, ACE richness increased with TOC (R = 0.23, p < 0.05), CDOM PC1 (R = 0.26, p < 0.03), α_ph_(λ); m^− 1^ (R = 0.25, p < 0.05) and a_CDOM_(400) (R = 0.27, p < 0.025), but not POC (for more details see Supplementary Figure S1). Further assessment of the associations by generalized additive models (GAMs) revealed that including non-monotonicity improved the models between alpha diversity (ACE richness and Shannon index), and organic matter descriptors considerably (i.e. as indicated by AIC, GCV, R^2^ and chisq; Supplementary Table S2). Resulting GAMs revealed a peak in alpha diversity (ACE richness and Shannon diversity) at intermediate browning, i.e. CDOM-PC1 and a_CDOM_ (Supplementary Figure S4). Such humped-shaped curves in association studies of alpha-diversity have been observed widely as for example when inferring relationships between productivity and biodiversity [2–4].

### Spatial and environmental factors affecting bacterioplankton community composition

Bacterial community dissimilarity increased significantly with geographic distance which, despite a pronounced scatter and low coefficient (R^2^ = 0.066), exhibited significant distance-decay relationships (p < 0.0001). Similarly, variance partitioning analysis revealed that the fraction of the total community variation that could be explained solely by spatial factors (longitude and latitude 1.2%) was small. In comparison, the fraction that could be solely explained by local environment conditions was 11.2% combined for CO2, TN, PO4 and temperature, and 5.0% for TOC, CDOM and a_CDOM_(400) while with shared effects 20.4% and 13.1%, respectively (Figure 2A). Approximately 72% of the community variance along the sampled lake gradient remained unexplained by the measured environmental and spatial gradient indicators assuming monotonic relationships.

**Figure 2.**
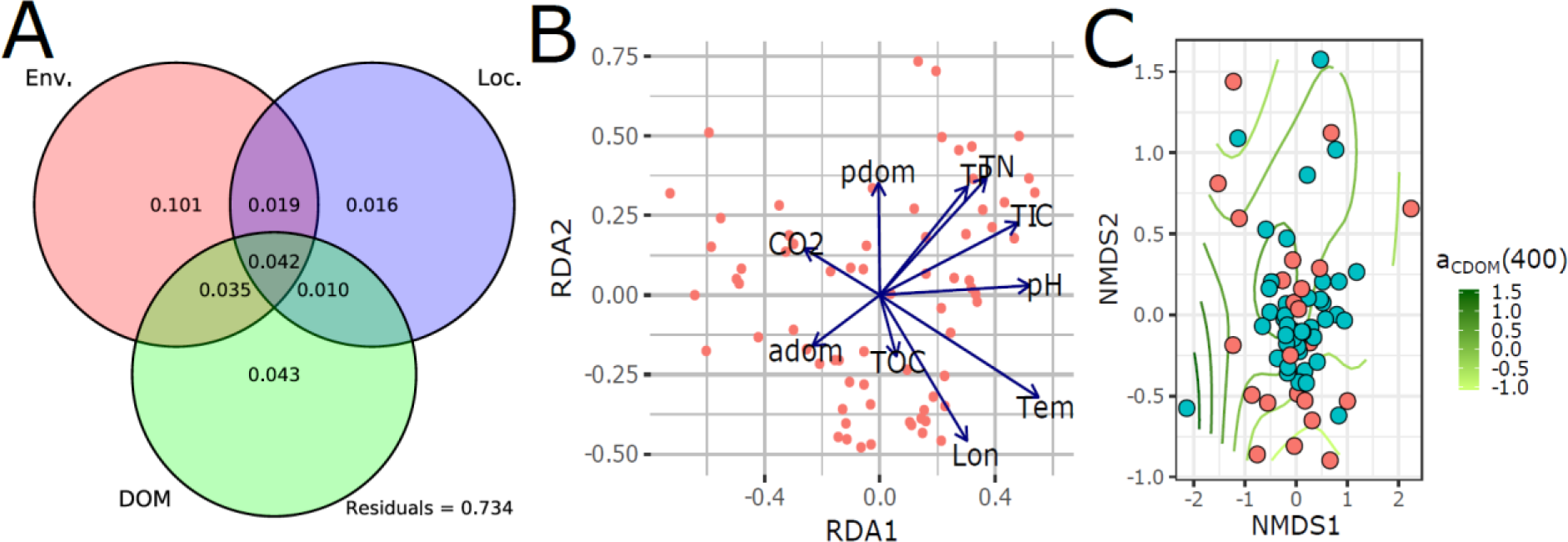
Partitioning of the total variance in the bacterial community **(A)** with environmental (Env.), organic matter properties (DOM) and spatial (Loc.) descriptors. Results from an unconstrained redundancy analysis **(B)** showing the covariation between the composition of bacterial communities and environmental factors. Arrows represent fitted gradient vectors for spatial (Lon – longitude) and environmental (Tem – water temperature, pH, CO2 – carbon dioxide, TOC – total organic carbon, TP – total phosphorus, TN – total nitrogen, TIC – total inorganic carbon, adom - a_CDOM_(400) a proxy for aromaticity of CDOM and pdom - absorption coefficient spectra of phytoplankton pigments) variables. Ordisurf **(C)** with a_CDOM_(400) revealing the non-monotonic relationship with bacterial community composition.

The environmental properties showing a high co-variance with bacterial community composition were both spatial and environmental gradients including longitude (R^2^ = 0.247, p = 0.001), latitude (R^2^ = 0.183, p = 0.001), temperature (R^2^ = 0.341, p = 0.001), and concentrations of total nitrogen (R^2^ = 0.242, p = 0.001), total phosphorus (R^2^ = 0.235, p = 0.001), CO2 (R^2^ = 0.174, p = 0.004) and PO4 (R^2^ = 0.384, p = 0.001). As revealed by RDA the direction of maximal increase for the fitted vectors representing longitude, temperature and CO2 was similar, but orthogonal to the vectors reflecting nutrient state (i.e. TN, TP and PO4 concentrations) (Figure 2B). This can be interpreted that there are two main directionalities driving bacterial community composition in lakes, corresponding to nutrient status and temperature.

Monotonic functions as used in RDA revealed short vectors for TOC, a_CDOM_(400) and CDOM-PC1 which can be interpreted that organic matter is a poor predictor of bacterial community compositions in lakes. However, there is no reason to assume that TOC, a_CDOM_(400) and CDOM-PC1 vary in a monotonic fashion across the RDA’s biplot (Figure 2B) which is a prerequisite to identify relationships in unconstrained ordination. To reveal potential non-monotonic relations, we fitted a smooth response surface of TOC and a_CDOM_(400) values over the limits of the biplot using *ordisurf* function (i.e. for a_CDOM_(400) see Figure 2C and for TOC Supplementary Figure S5). The fitted surfaces are far from monotonic and revealed that the relationships of a_CDOM_(400) and TOC with the bacterial community are significant (p < 0.001) and explained large parts of the variability (a_CDOM_(400): adj. R-square = 0.3; deviance explained 41.8%; and TOC: adj. R-square = 0.30; deviance explained 36.5%) when performing smoothness selection via Marginal (Maximum) Likelihood model fitting.

Similar beta diversity patterns appeared along the a_CDOM_(400) gradient for both XGBoost, random forest and FFNN models (Figure 3A-C). The mean value of the response surface (i.e. 0.916 in the XGBoost models for TOC and a_CDOM_(400)) can be treated as the baseline beta diversity across all sites. Data points with values below the mean present higher similarity between sites; likewise, higher values represent lower similarity. Data points located on the diagonal are not presented as they are pairwise distances of a site to itself, thus assumed to be zero. To interpret the response surfaces, one may begin by looking at a point bordering the diagonal and then follow a line of points further up on the a_CDOM_(400) site 2 axis. A “ridge” indicates a a_CDOM_(400) value next to the diagonal to be a likely threshold from which the shift in bacterial community composition is greater than average. In the same manner, a “valley” indicates a a_CDOM_(400) value next to the diagonal is likely located on an interval of the a_CDOM_(400) gradient along which bacterial communities do not shift substantially. Following this interpretation, a_CDOM_(400) thresholds for high variation in BCC appear around 0.3, 0.5 and 1-1.5 absorbance units, while communities are more similar to others with higher a_CDOM_(400) around 0.4, 0.6 and 1.6-2.25 absorbance units. In comparison, the linear model captured the greater variation in beta diversity pattern above 2 absorbance units on the a_CDOM_(400) gradient (Supplementary Figure S6), but not the multiple ridges or valleys revealed by XGBoost, random forest regression and FFNN (Figure 3A-C).

**Figure 3.**
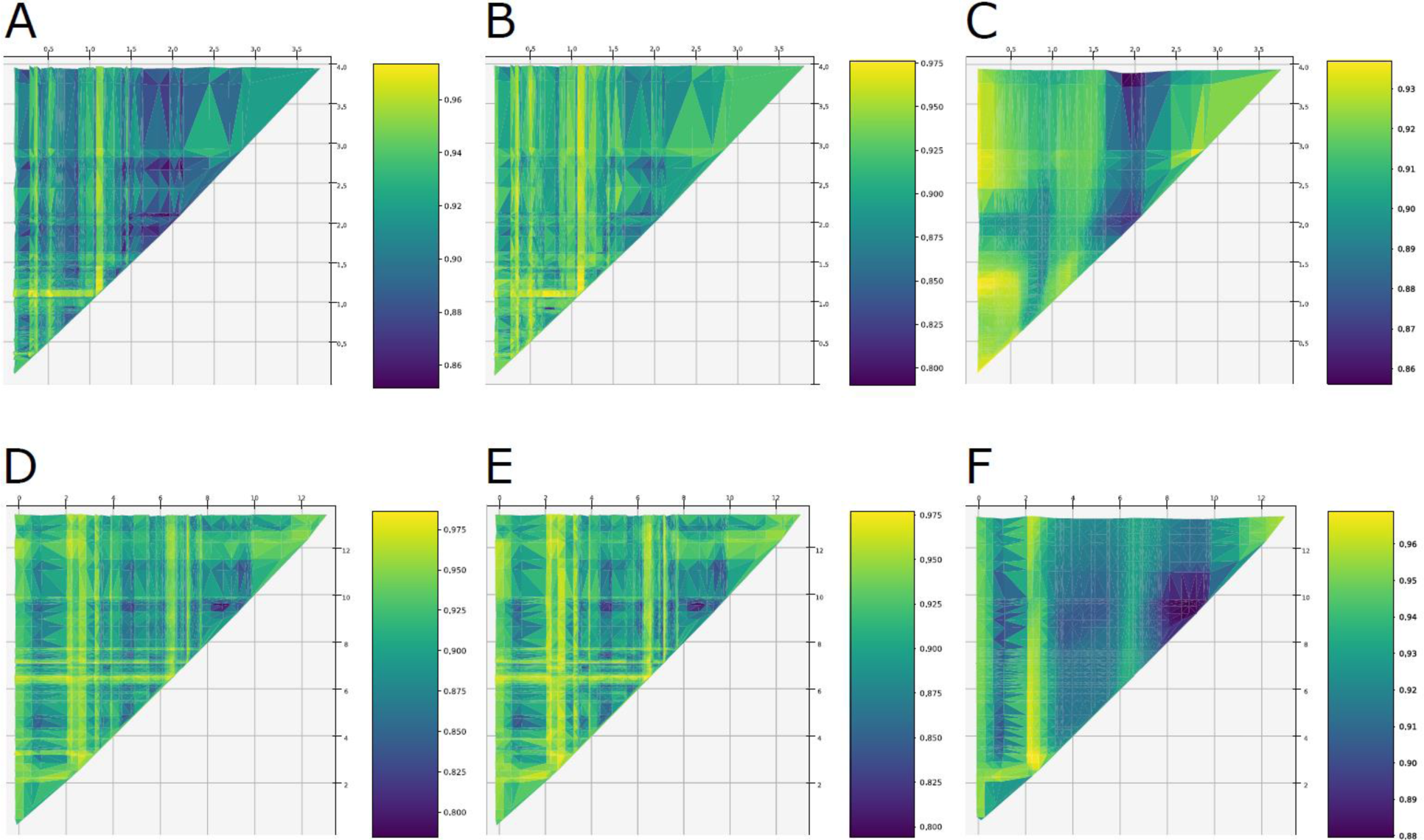
Visualization of XGBoost (**A, D**), random forest (B, E) and feed-forward neural network (C, F) predictions of bacterial community compositional changes (Bray-Curtis distances). Compositional changes were predicted for a meshgrid of a_CDOM_(400) **(A-C)** and TOC **(D-F)** values spanning the minimum-maximum range of the gradient with a step size equal to the smallest pairwise a_CDOM_(400) and TOC differences. The mean value of the response surface can be treated as the baseline beta diversity across all sites. Data points with values below the mean represent higher similarity between sites; likewise, higher values represent lower similarity. To interpret the response surfaces, one may begin by looking at a point bordering the diagonal and then follow a line of points further up on the a_CDOM_(400) or TOC site 2 axis. Here, a “ridge” indicates a a_CDOM_(400) or TOC value next to the diagonal to be a likely threshold from which the shift in bacterial community composition is greater than average. In the same manner, a “valley” indicates a a_CDOM_(400) or TOC value next to the diagonal which is likely located on an interval of the a_CDOM_(400) or TOC gradient along which bacterial communities do not shift substantially.

Similarly, model results revealed “ridges” along the TOC gradient (Figure 3D-F), indicative for thresholds at which shifts in BCC are greater than average. These TOC thresholds for high variation in BCC appear around 0.3, and 2-3 and 6.5 mgC L^-1^. “Valleys” indicative for an interval of the TOC gradient where bacterial communities do not shift substantially were predicted to be around 1.5, 4-5 and 8 mgC L^-1^. Overall, R^2^ values of the XGBoost model predictions (R^2^TOC = 0.446; R^2^a_CDOM_ = 0.315) showed smaller differences between the observed data and the fitted values, than the FFNN (R^2^TOC = 0.068; R^2^a_CDOM_ = 0.014) and the random forest (R^2^TOC = 0.414; R^2^a_CDOM_ = 0.172) models. Training the models with and without the duplicate samples did not affect the models.

### Association between DOM and bacterial taxonomic groups

Altogether, 29 bacterial phyla were detected resembling results in line with the global synoptic meta-analysis of 16S rRNA gene sequences from lake epilimnia [57] and Zwart *et al.* [58], showing that 4 phyla (Proteobacteria, Actinobacteria, Bacteroidetes and Cyanobacteria) were recovered commonly across the sampled freshwater ecosystems (Supplementary Figure S7A). With regards to number of ASVs, Proteobacteria was the most diverse phylum (4414 ASVs) followed by Bacteroidetes (1317 ASVs), while Cyanobacteria and Actinobacteria had similar richness (784 and 652 ASVs, respectively) (Supplementary Figure S7B). By further resolving the taxonomy to the genus level, the most abundant identified groups were alfIV-A (LD12) (6.0%), *Aquinocola* (4.9%), various acI (4.6%), *Synechococcus* (3.1%), *Niveitalea* (2.1%) and *Methyloferula* (1.5%).

To explore relationships between ASVs and environmental properties, we used MINE [56]. While this nonparametric approach identifies relationships of ASVs with all measured environmental variables, we will focus on the results from the analyses with a_CDOM_(400) and TOC. MINE identified 108 significant relationships (q-value < 0.05) with a MIC of 0.316 to 1 between single ASVs and TOC while 92 ASVs were identified with significant relationships with a_CDOM_(400) with a MIC of 0.316 to 1. The maximum asymmetry score (MAS) ranged from around 0.05 to 0.58 for a_CDOM_(400) and 0.05 to 0.67 for TOC. MAS values below 0.05 indicate a monotone relationship between ASVs and a_CDOM_(400) or TOC (Figure 4). While purely monotone relationships were not detected, non-monotonic responses dominated which are indicative of the existence of thresholds in the response of ASVs along the sampled CDOM and TOC gradients, similar to the model predictions of the entire bacterial community responses.

**Figure 4.**
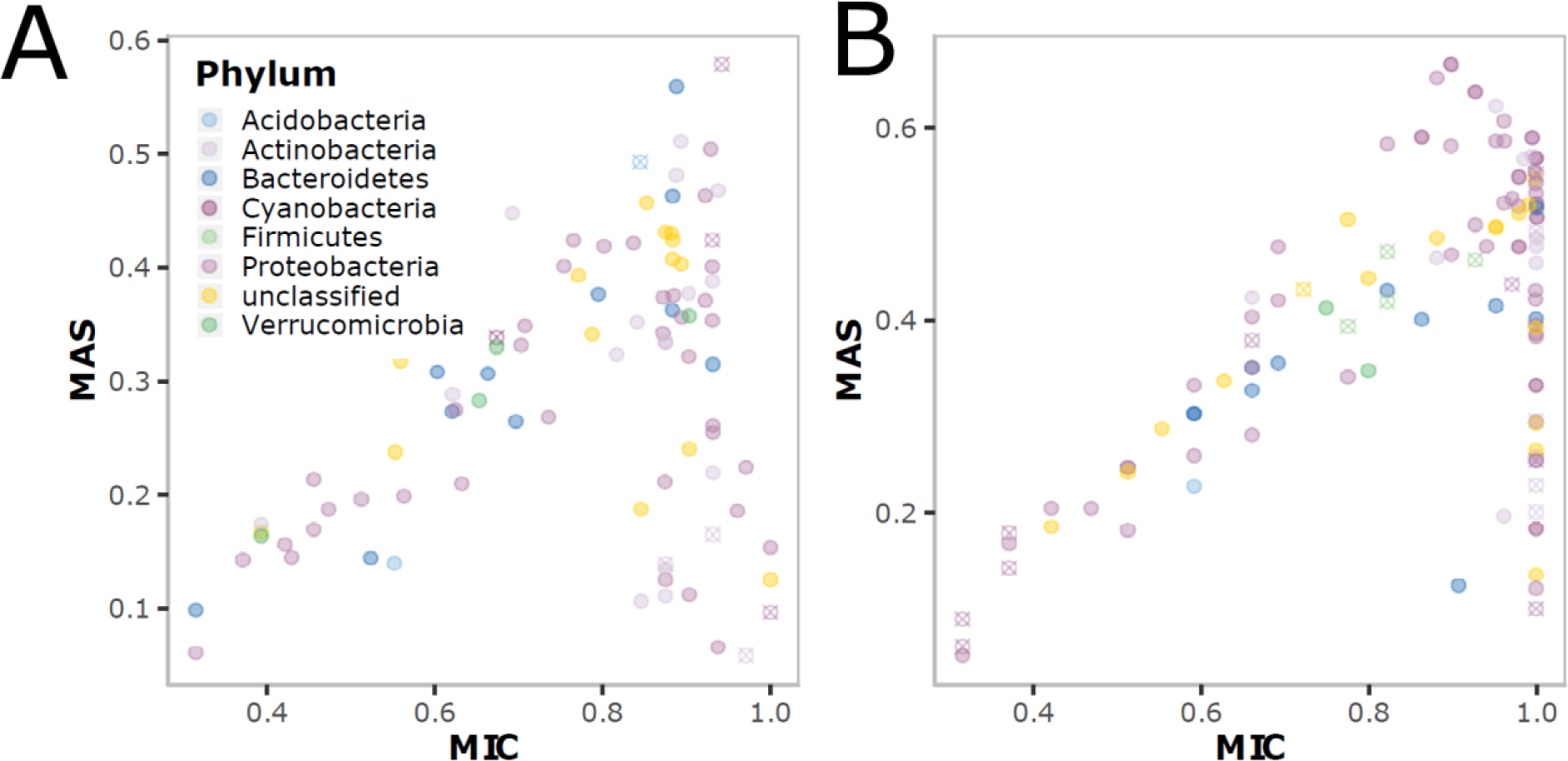
Plots summarizing MINE statistics of the relationships between ASVs and a_CDOM_(400) **(A)**, and ASVs and TOC **(B)**. Depicted MINE statistics are MIC - coefficient, MAS – non-monotonicity. The color of the symbols indicates the taxonomic affiliation of the ASV at the phylum level.

## Discussion

Our longitudinal survey showed that monotonic functions are poor predictors of the relationship between browning and microbial diversity. Both alpha and beta diversity were poorly predicted by monotonic functions, as the variation explained was scarcely exceeding 5 % when using linear models, RDA and variance partitioning. The variation explained increased with models taking deviations from monotonicity into account. For example, the fraction of variance explained in beta diversity increased up to 45 % when using XGBoost, 41 % with random forest regressor and 6.8 % with FFNN while 30% with Marginal Likelihood models. In addition, we demonstrate that most relationships between bacterial taxa (ASVs), and TOC concentrations and chromophoric properties of the water were non-monotonic. A common feature of non-monotonic functions is that they define relationships with both increasing and decreasing sectors as well as different stable states (“valleys”) where the nature of the response can change suddenly when an environmental factor (i.e. browning) reaches a threshold (“ridge”). Results from the Marginal Likelihood model fitting can be interpreted along these lines since the model reveals distinct a_CDOM_(400) and TOC types coinciding with distinct environmental conditions and bacterial community composition profiles. Such non-monotonicity in response to DOM (a complex of substrates for microbial growth) can be predicted from kinetics studies emphasizing that growth may not be controlled by only a single compound but by two or more compounds simultaneously, and that kinetic properties of a community might change due to adaptation of individual cells or community composition to ever changing environmental conditions [59].

To capture further details and validate the findings of the Marginal Likelihood model fitting such as thresholds and non-monotonicity in bacterial community responses along the DOM gradient, we applied machine learning methods, in particular FFNN, random forest and XGBoost.

A key finding revealed by the machine learning methods is the apparent presence of multiple thresholds (“ridges”) along the a_CDOM_(400) and TOC gradients where bacterial community composition shifts, corroborating the predictions of the Marginal Likelihood model fitting. If bacterial community composition had been found to vary linearly along the TOC and a_CDOM_(400) gradients, they would have presented a pattern of isolines parallel to the diagonal in Figure 3; and there would have been a linear relationship between BCC distances and a_CDOM_(400) or TOC differences between sites (Supplementary Figure S6). The observed thresholds (“ridges”) in TOC concentrations and a_CDOM_(400) values can be interpreted as “guardrails” of biodiversity along the browning gradient. These “guardrails” guard alternative trajectories at low browning which converge into a single almost monotonic (linear) trajectory when TOC concentrations (above 10 mgC L^-1^) or chromophoric properties reach high levels (above 2.5 absorbance units). At low TOC and a_CDOM_(400), community patterns can be interpreted as resembling a phenomenon known as hysteresis where multiple states persist under equal environmental conditions. However, in contrast to alternative stable state theory which suggests discrete states separated by ecological thresholds, our results point instead to alternative trajectories in biodiversity separated by “guardrails” which start at 0.3, 0.5 and 1-1.5 absorbance units of a_CDOM_(400), respectively. For TOC the separating “guardrails” are predicted to start at around 0.5, and 2-3 and 6 mgC L^-1^.

Comparisons of the three applied machine learning approaches revealed similar beta diversity patterns along the a_CDOM_(400) and TOC gradients. XGBoost reflected the raw data more closely (greater R^2^) and was orders of magnitude faster than the FFNN. This corroborates previous observation that decision tree based algorithms such as XGBoost outperform neural networks when small-to-medium structured/tabular data is used, as in our case.

A step in the causal chain to explain the apparent non-monotonicity in the relationship between browning and biodiversity is the non-interchangeable nature of individual taxa responses. Individual taxa responses can be direct and indirect with opposite and non-additive strategies based on changes in the environment. This is reflected by browning mostly leading to non-monotonic relationships as shown by the high number of ASVs with high MAS (figure 4). The non-monotonicity in response to environmental stimuli can be explained by organisms’ ability to adopt opposite strategies along the stimulís gradient. In the case of browning terrestrially derived TOC provides a significant source of C for heterotrophic bacteria [8, 60] and where different fractions of this TOC are utilized with different efficiency [61]. The different fractions are also utilized by different taxa, which, as shown in our study, leads to different ASVs being present along the browning gradient. These opposing positive and negative effects on individual ASVs are only monotonic if they change in the same order or scale, so that their net effect will be additive. However, if the positive and negative effects change in different orders or scales, which is common in nature, their net effect will not be additive, and the function will be non-monotonic. This is reflected in the high number of non-monotonic relationships in the co-occurrence patterns among ASVs (Supplementary Figure S8). Additional potential explanations for the apparent non-monotonic responses of individual taxa are shifts in interaction behavior with examples such as the Prisoner’s Dilemma [62] and opposing dual effects between organisms.

As shown by previous studies, seasonality, water mixing as well as source and age of TOC clearly offer different sources of energy that may select for different microbial community members and metabolic pathways at both short and long timescales. The non-monotonic responses in community composition, as observed in our study, are likely also reflecting a tradeoff between nutrients associated with CDOM and the increasing light attenuation caused by CDOM. Modest increases in TOC and CDOM have been shown to block out short-wave UV-radiation [63] and to limit autochthonous production of TOC. Since browning is increasing by processes associated with climate change [11, 13] and the strong decline of atmospheric sulfur (S) deposition [12, 13], we predict, by translating our model results based on spatial data into a temporal context, that lake bacterioplankton diversity will develop along different trajectories (“valleys”) guided by thresholds (“ridges” or “guardrails”) at low browning (i.e. low TOC concentrations and low chromophoric properties). Lakes with high levels are predicted to follow a closely monotonic trajectory of biodiversity change over time. Considering that browning is an ongoing process, alternative trends of bacterial diversity in lakes currently experiencing low TOC and CDOM levels are expected while more uniform and monotonic trends are predicted in lakes with high levels of browning (above 10 mgC L^-1^).

To conclude, comparing machine learning results, in particular XGBoost, with the outputs of the regressions and RDA model revealed how poorly linear models perform when attempting to capture the association between browning and microbial biodiversity. In addition to poor predictive power linear models will miss idiosyncratic patterns in biodiversity as they will be overwritten and clumped by monotonic functions. As such our results highlight the need to explore non-monotonic relationships common in biological systems which might provide part of the explanation of contrasting results among different studies, in addition to revealing the real complexity of associations between biodiversity and environmental properties. Most importantly, by using non-monotonic functions and models the position of thresholds, alternative trajectories and guardrails can be revealed which are important for mitigation efforts and management decisions to counteract environmental changes not only in freshwater microbiomes affected by browning.

## Acknowledgements

This study has been supported financially by the Department of Biosciences and the Centre of Biogeochemistry in the Anthropocene, University of Oslo, and by the Research Council of Norway (grant ‘ECCO’ 224779 to Dag O. Hessen and grant ‘COMSAT’ 196336 to Tom Andersen). We thank the COMSAT field sampling crew, especially Johnny Håll, Marcia Kyle, Robert Ptacnik, and Jan-Erik Thrane, for their efforts. Berit Kaasa and Sissel Brubak are acknowledged for laboratory support.

## Contribution

This study was designed by TA and DOH. Sample and data collection were coordinated by TA. Molecular analyses were performed by MK under supervision of SR and MD. Data was analyzed and visualized by MK, LF, SR, TA and AE. The first version of the manuscript was drafted by MK but substantially modified after additional data analysis by LF and AE. All authors provided comments and were involved in writing the final version of the manuscript. Financial support for the project was acquired by TA, DOH and AE.

## Competing interests

The authors declare no competing interests.

---

**Supplementary figure S1.**
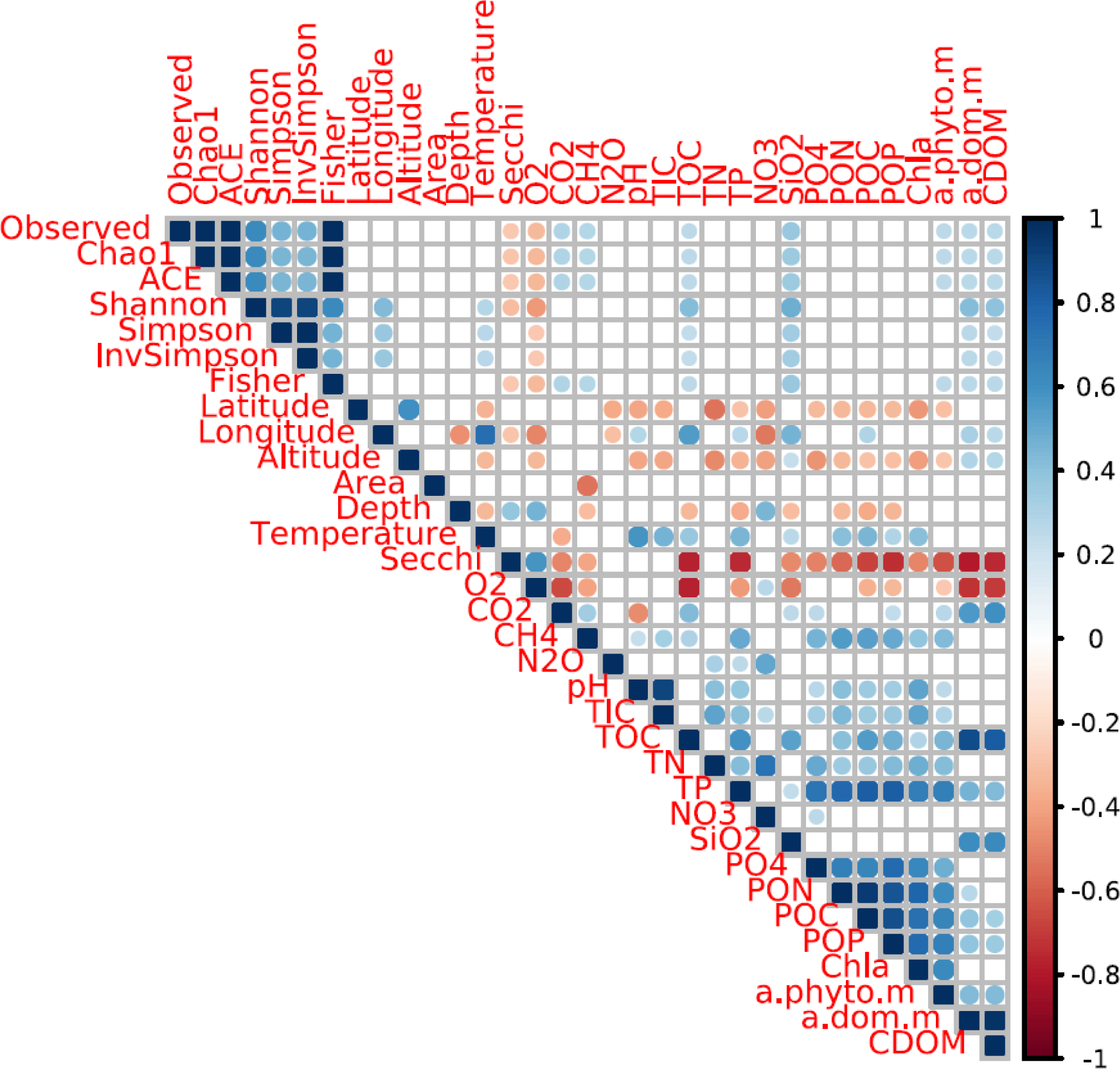
Cross-correlation statistics between environmental properties and alpha diversity (richness and evenness) measures. The size of the symbols corresponds to the significance level (larger symbols indicate smaller p value) whereas the color corresponds to the correlation coefficient (R).

**Supplementary figure S2.**
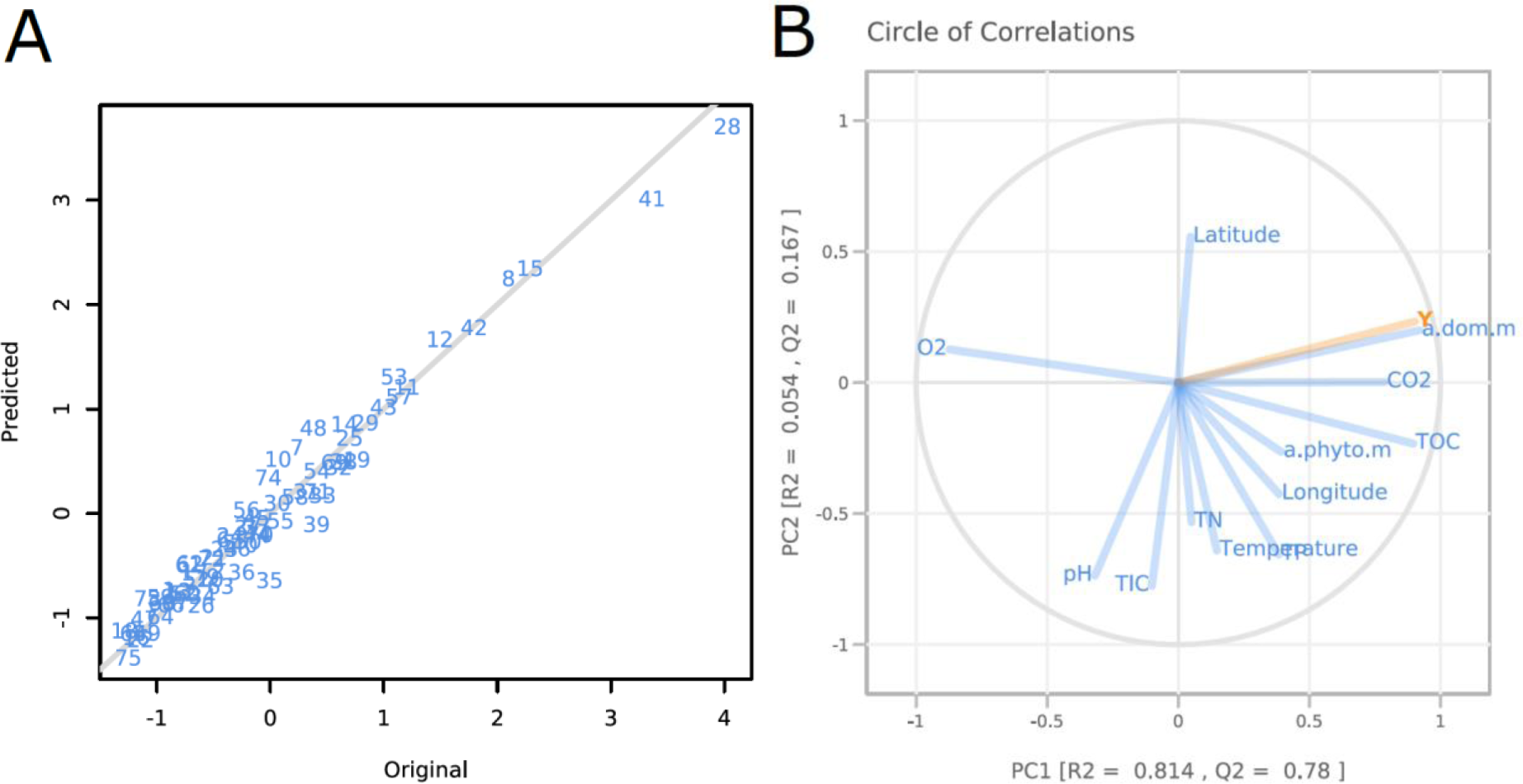
Comparison of observed and partial least square (PLS) model predictions of CDOM-PC1 (Principal component 1 describing 88% of the variability in fluorescence spectra of dissolved organic matter among the studied lake systems) corroborating the high predictive power of the PLS model when using environmental properties **(A)**. In panel **B** a scoring plot of an alternative PLS model including the first principal component (describing 88% of the variability in fluorescence spectra of dissolved organic matter among the studied lake systems) is shown.

**Supplementary figure S3.**
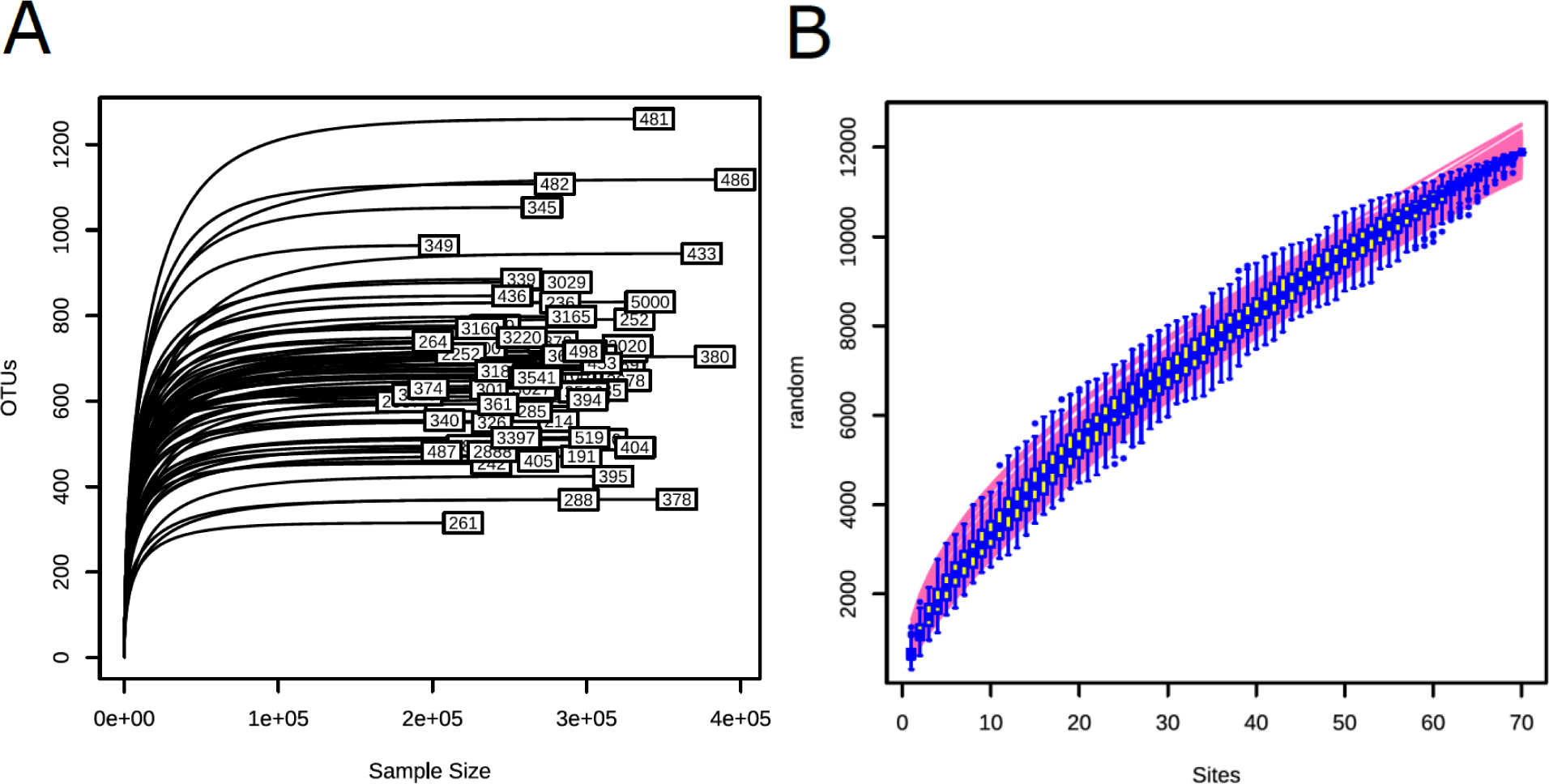
Rarefaction curves of bacterial sequence variants (ASVs) richness for each lake **(A)** and the sampled region (all lakes combined; **B**).

**Supplementary figure S4.**
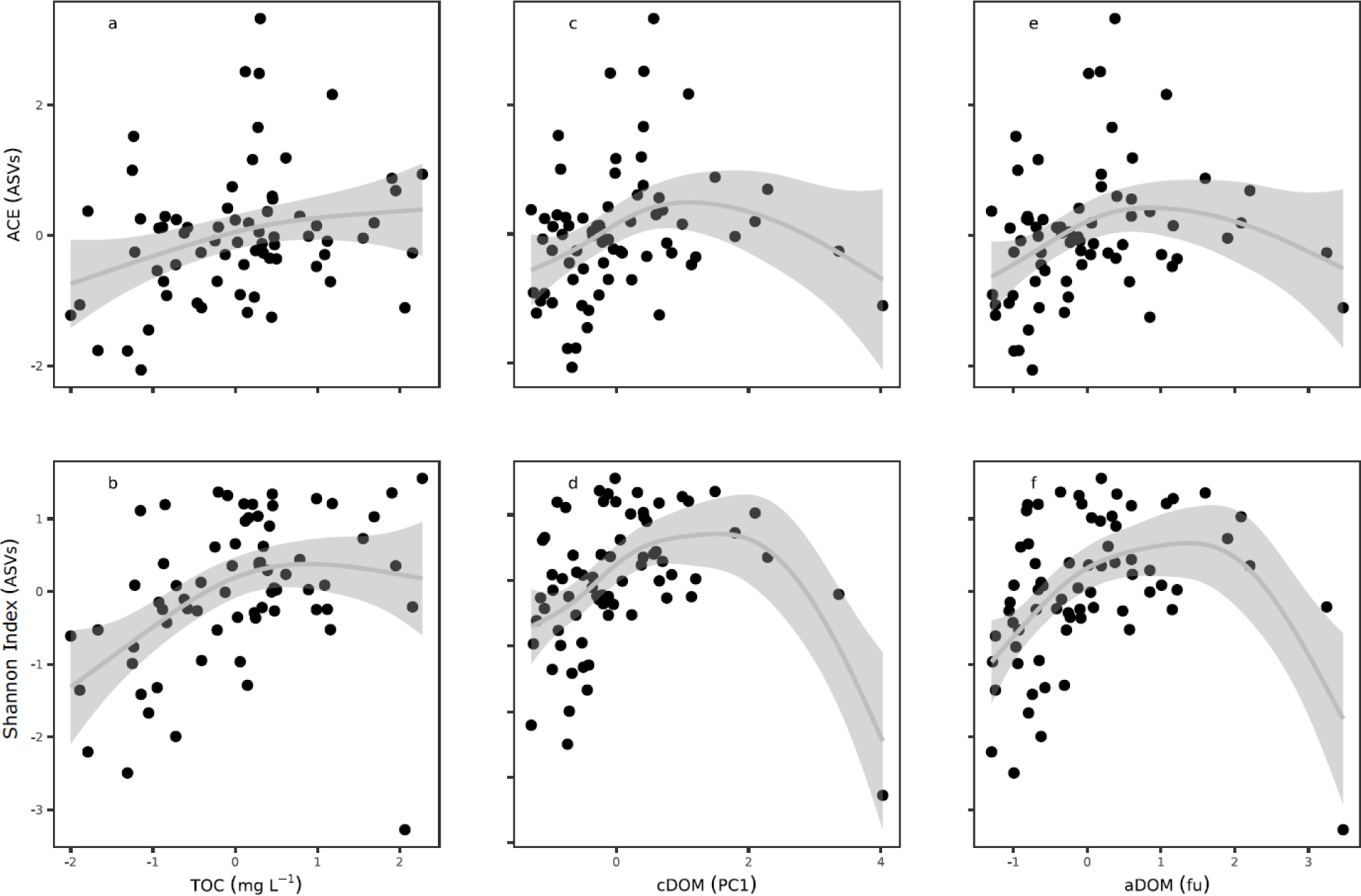
Generalized additive model plots revealing the non-monotonic association between alpha diversity measures (ACE and Shannon index) and total organic matter (TOC; **a, b**), chromophoric dissolved organic matter (cDOM; **c, d**) and aromaticity of CDOM (aDOM; **e, f**). Model statistics and comparisons with linear models are given in Supplementary Table S2.

**Supplementary figure S5.**
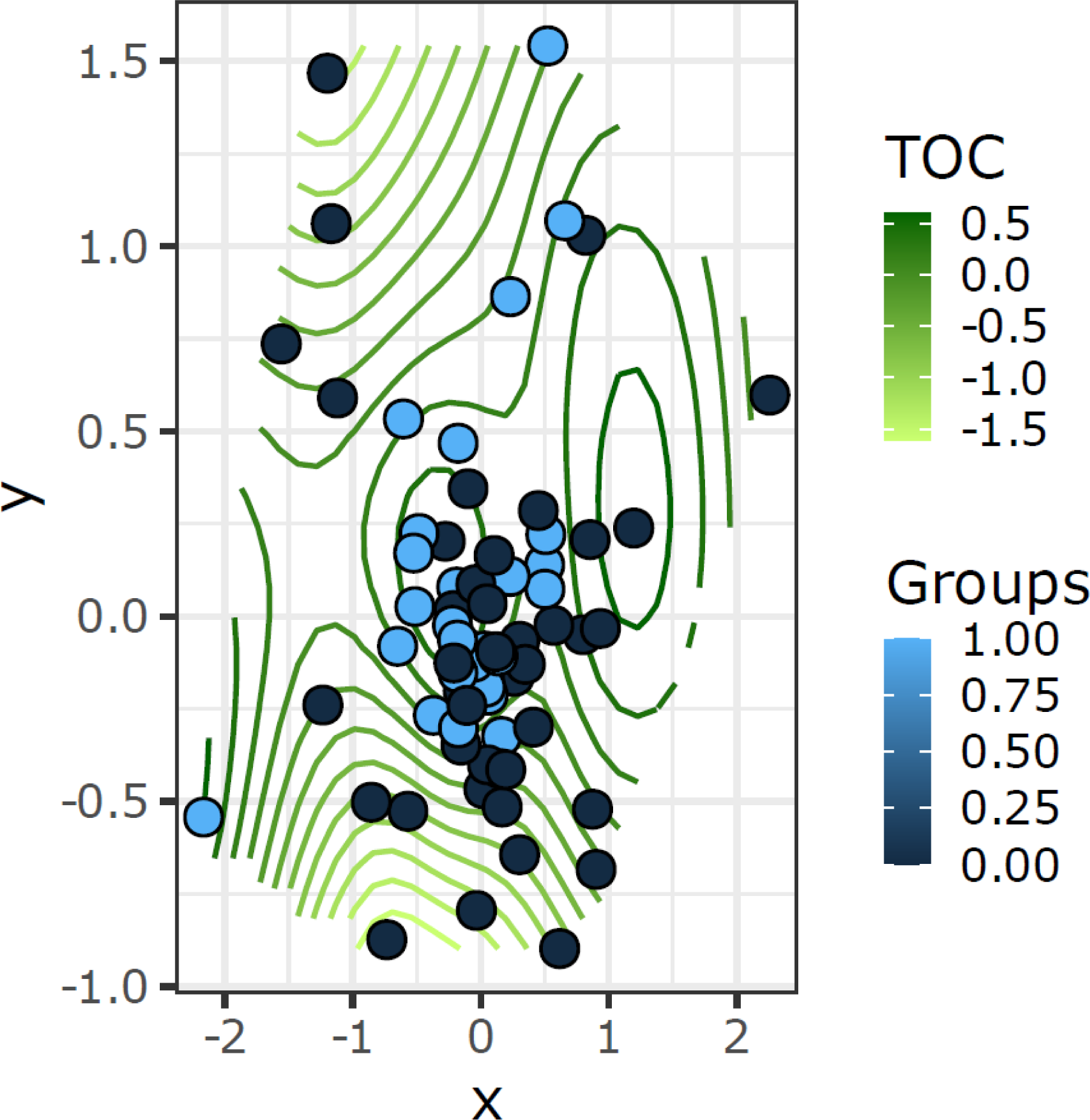
Ordisurf contour plot for total organic carbon (TOC) illustrating the non-monotonic relationship with bacterial community composition.

**Supplementary figure S6.**
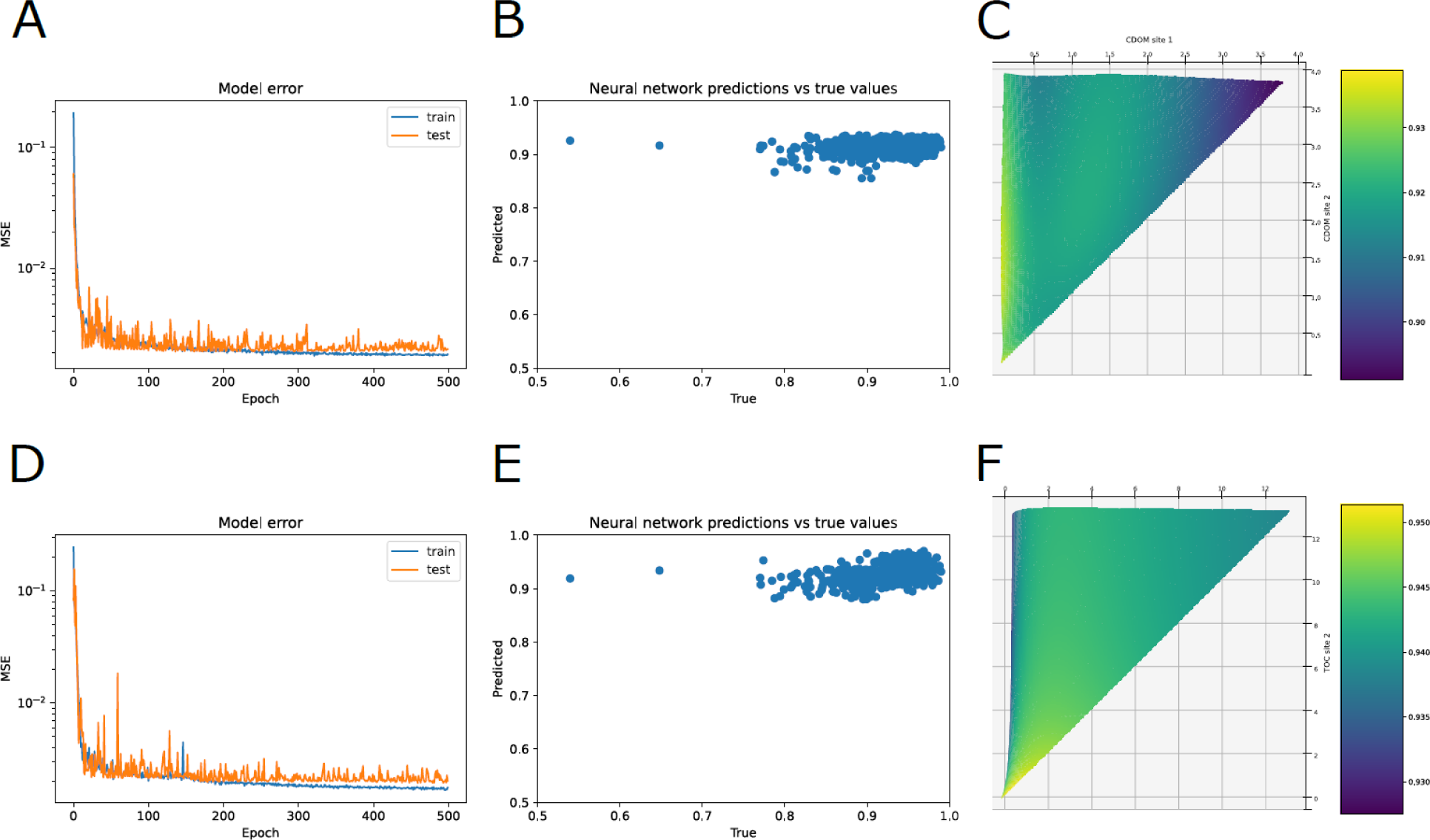
Validation plot of the neural networks shown in figure 3. Mean square errors (MSE) of training, validation and testing for predicting bacterial compositional changes from a_CDOM_(400) **(A)** and TOC **(B)**. Raw versus predicted values are plotted in **C** for a_CDOM_(400) and in **D** for TOC. Linear models are given in **E** for a_CDOM_(400) and in **F** for TOC .

**Supplementary figure S7.**
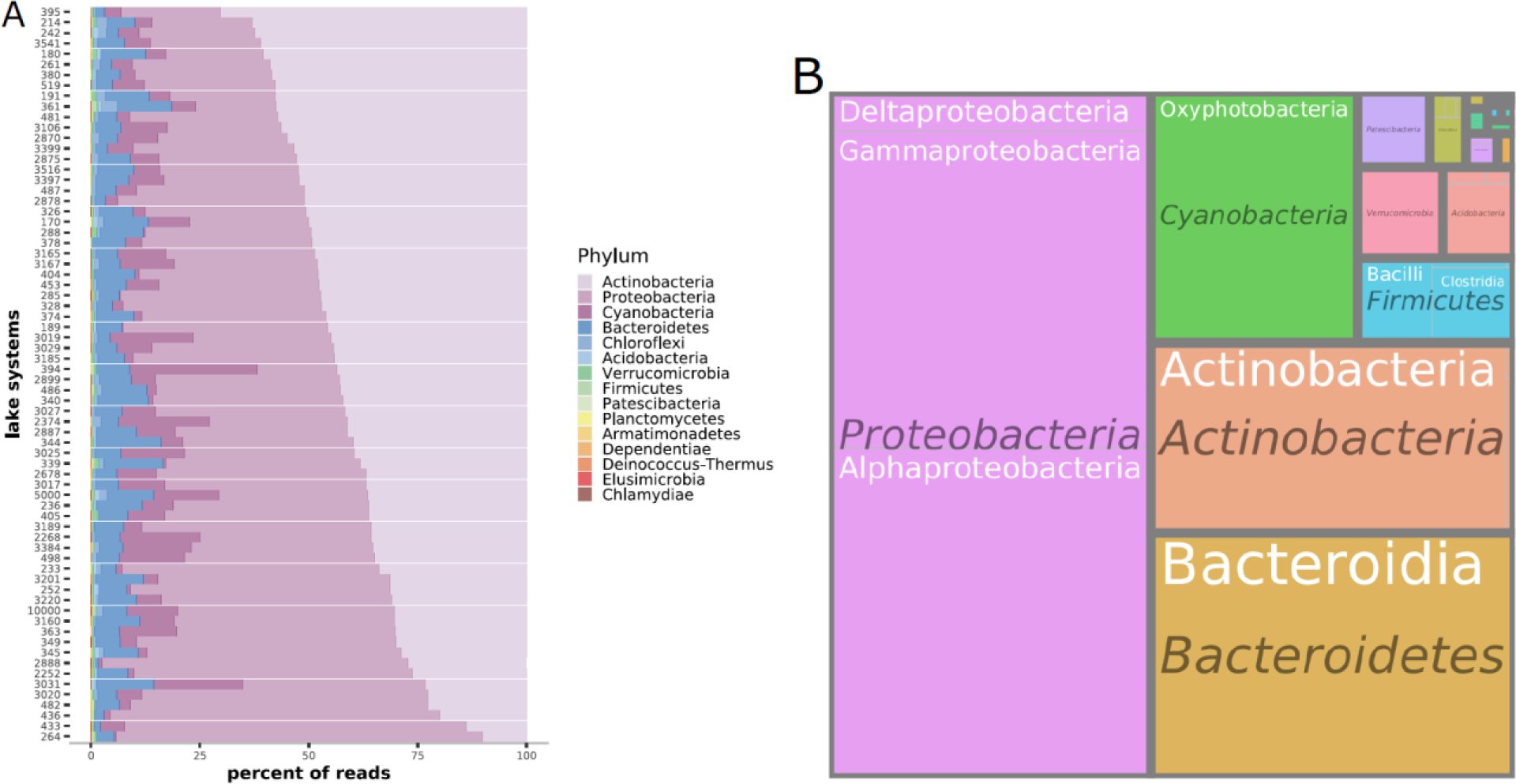
Treemap plots showing the proportion of different phyla (and most common classes) in regards to the number of reads **(A)** and bacterial sequence variances (ASVs; **B**). These Treemaps display hierarchical (taxonomic) data as a set of nested rectangles. Each branch (Phylum) of the taxonomy tree is given a rectangle, which is then tiled with smaller rectangles representing sub-branches (classes). A leaf node’s rectangle has an area proportional to a specified dimension of the data – proportion of reads **(A)** and proportion of ASVs.

**Supplementary figure S8.**
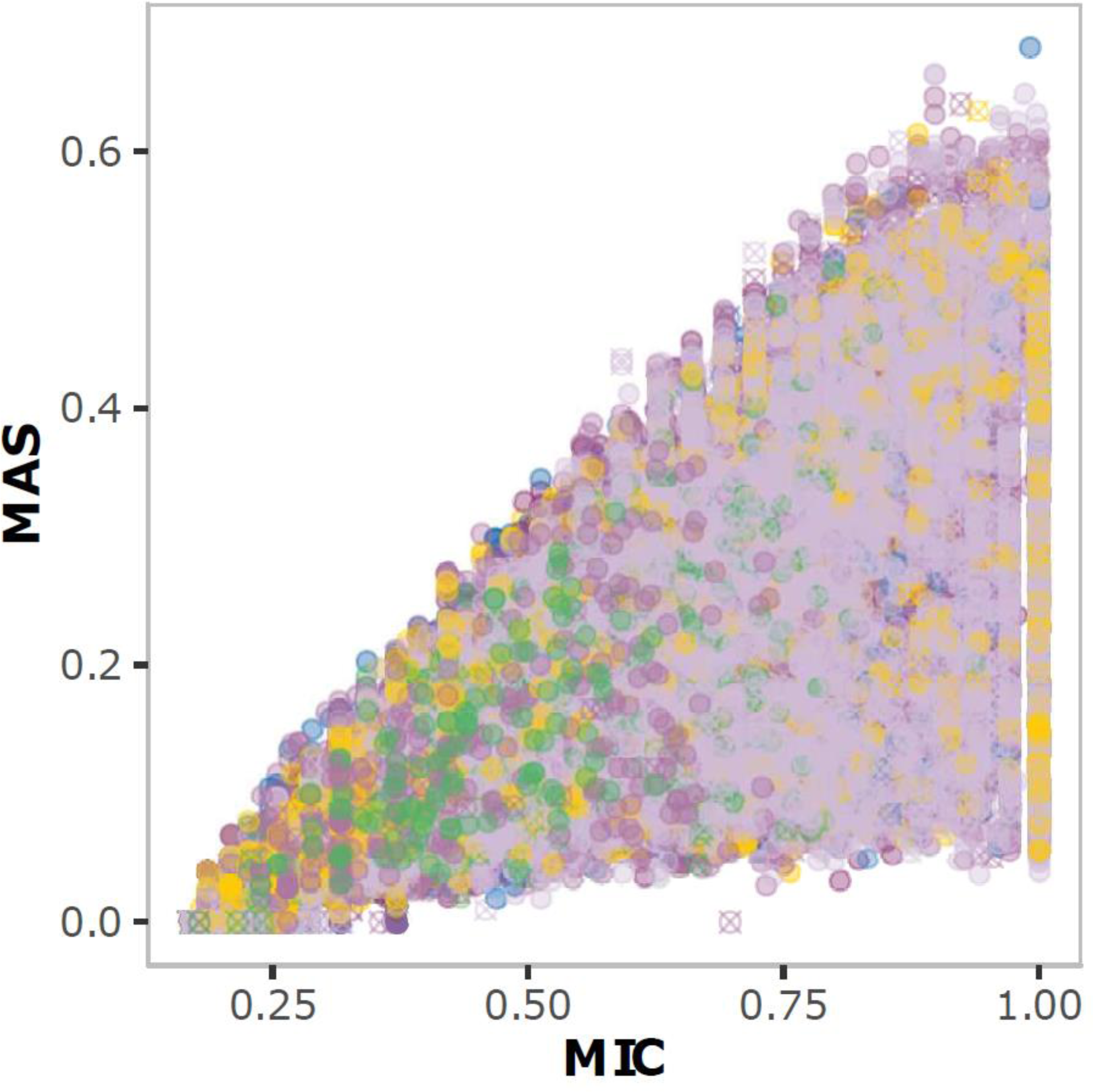
Plot summarizing MINE statistics of the relationships between ASVs. Depicted MINE statistics are MIC – coefficient, MAS – non-monotonicity. The color of the symbols indicates the taxonomic affiliation of the ASV at the phylum level.

**Supplementary Table S1.**
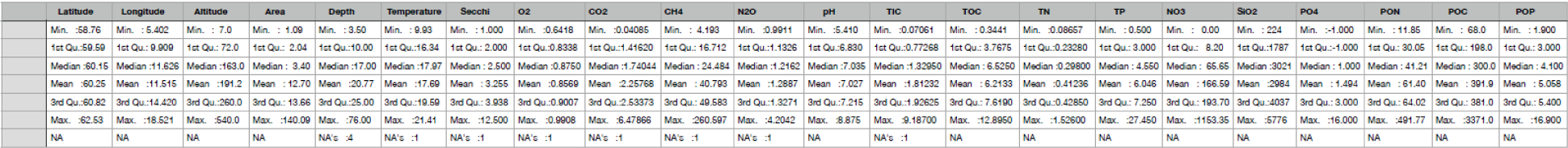
Complete list of environmental variables and summary statistics.

**Supplementary Table S2.**
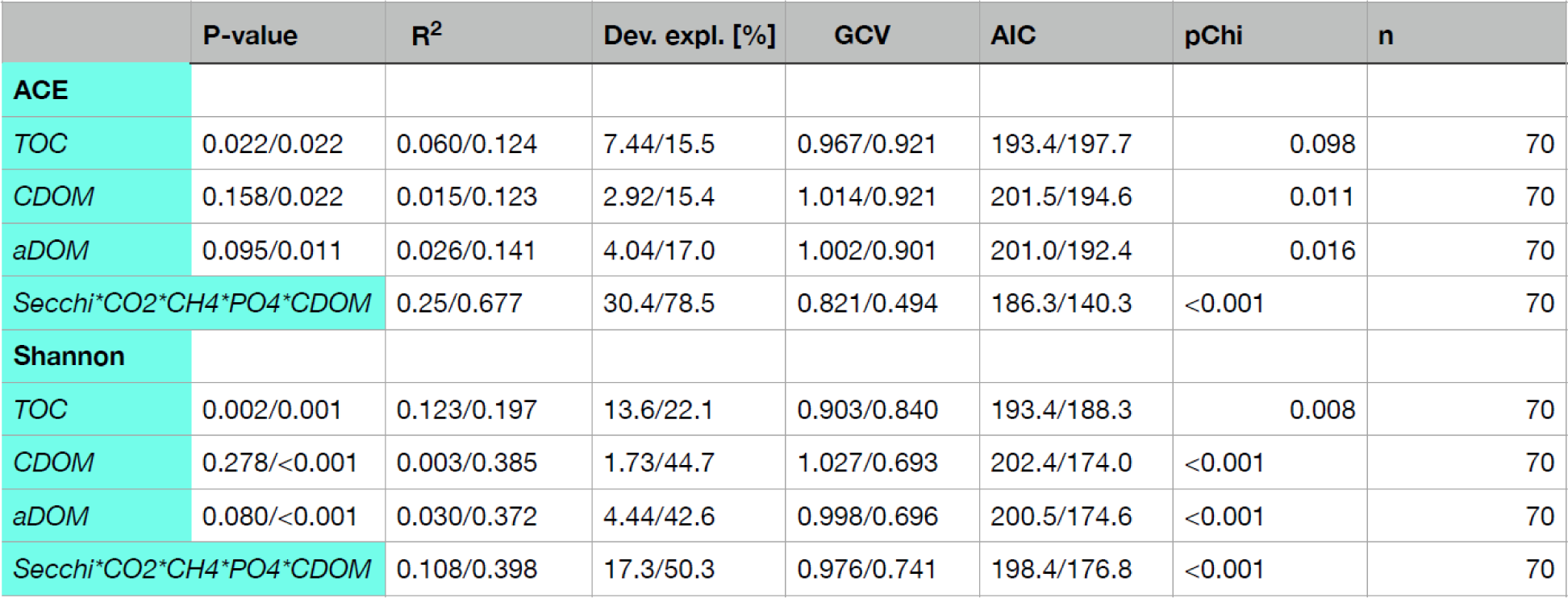
Statistical comparison of linear and generalized additive models as given by the p-value, deviation explained (Dev. expl.), Akaike information criterion (AIC), generalized cross-validation (GCV), coefficient of determination (R^2^), the p-value of the Chi-squared test (pChi) and the number of samples (n).

## Notes

https://www.ncbi.nlm.nih.gov/bioproject/?term=PRJNA637765

### Competing Interest Statement

The authors have declared no competing interest.

